# BCG immunization induced KLRG1+ NK cells show memory-like responses to mycobacterial and HIV antigens

**DOI:** 10.1101/2024.05.09.593411

**Authors:** Manuja Gunasena, Mario Alles, Thorsten Demberg, Will Mulhern, Namal P.M. Liyanage

## Abstract

The live-attenuated Bacille-Calmette-Guérin (BCG) vaccine is the only approved vaccine against *Mycobacterium tuberculosis* (MTB), offering broad protection against tuberculosis (TB) and other infectious diseases. ‘Trained immunity’, a process where innate immune cells develop memory-like features, is considered one of the BCG vaccine’s protective mechanisms. In this study, we investigated the effect of BCG vaccination on Natural Killer (NK) cells, a key subset of the innate immune system, and their ability to give rise to heterologous memory-like responses to HIV antigens. Here we found that BCG vaccine-induced KLRG1+ NK cells exhibit memory-like responses to both MTB and HIV antigens, as evidenced by their increased production of IFNγ upon exposure to MTB and HIV-gag antigens. This finding is of great importance, as co-infection with HIV and TB is highly relevant in Asia and Africa where BCG is administered. Understanding these responses is crucial for the development of more effective vaccines and therapeutics for both TB and HIV.

## Introduction

The live-attenuated BCG vaccine is the only approved vaccine against MTB (1, 2). BCG is typically administered intradermally at birth or soon after and is mainly protective against tuberculous meningitis and miliary tuberculosis in infancy and childhood(3). BCG vaccination not only protects against TB in children but also lowers childhood mortality from other infectious diseases, particularly in Southeast Asia and Africa (4). Additionally, BCG vaccination offers broad off-target protection against several pathogens, although its precise mechanism of action remains unknown. ‘Trained immunity’, a process where innate immune cells develop memory-like features, is thought to be one of the BCG vaccine’s protective mechanisms. NK cells, a key subset of the innate immune compartment, have been shown to be modulated by BCG, exhibiting enhanced responses not only to homologues (*Mycobacterium*) antigens but also to heterologous (*Candida albicans* and *Staphylococcus aureus)* antigens (5).

The recall responses of innate immune cells are mediated through epigenetic and metabolic reprogramming (6). This reprogramming can be triggered by a wide range of stimuli, including BCG (7-10). Studies have shown that both human and murine NK cells can exhibit memory-like phenotypes, as demonstrated by their increased production of IFNγ upon re-stimulation (11, 12). These memory-like responses are associated with underlying epigenetic alterations, including reduced suppression of DNA methylation in the IFNγ gene locus and increased accessibility of genes regulating cell activation for transcription. This allows for a faster response upon re-stimulation(13). Moreover, studies have revealed that only certain phenotypes of NK cells, expanded following exposure to a training stimulus, undergo epigenetic remodeling in specific chromosomal regions. These regions, such as the *IFNG* conserved non-coding sequence, enable enhanced IFNγ production in these cells following re-exposure (13).

In this study, we investigate the effect of BCG-induced memory-like NK cells on both mycobacterial and HIV antigens. Here, we found that BCG vaccine-induced KLRG1+ NK cells exhibit memory-like responses to both mycobacterial antigens and HIV antigens, as evidenced by their increased production of IFNγ upon exposure to mycobacterial and HIV-gag antigens. To our knowledge, this is the first demonstration of BCG-induced memory-like NK cell responses to HIV antigens. This finding is of great importance as co-infection with HIV and TB is highly relevant in Asia and Africa where BCG is administered. Understanding these responses is crucial for the development of more effective vaccines and therapeutics for both TB and HIV.

## Material and Methods

### Ethics Statement

The study received approval from The Ohio State University Institutional Biosafety Committee and Institutional Animal Care and Use Committee, The study was performed in accordance with relevant institutional guidelines and is reported in accordance with ARRIVE guidelines (https://arriveguidelines.org).

### BCG Culture

*Mycobacterium bovis* BCG Pasteur strain cultures were prepared as per established protocols(14), including cultivation in media supplemented with OADC and Tween 80.

### Mouse Strains

C57BL/6 mice from The Jackson Laboratory were housed in controlled conditions with ad libitum access to food and water.

### Immunization and Tissue Collection

Mice were intramuscularly immunized with 1×10^4^CFU of BCG (i.e.,200μL of a 5×10^4^ CFU/mL aliquot) into the left thigh muscle (quadriceps), and spleens were collected on post-immunization day 18 for cell isolation.

### Cell Isolation

Spleens were pressed through a mesh screen using a syringe plunger, and cell suspensions were filtered through a 100-micron filter into a 15mL tube with ice-cold R10 media. After centrifugation, the cell pellet was resuspended, washed, and subjected to RBC lysis buffer before being spun down again at 250xg(14).

### Preparing *Mycobacterium bovis* Peptide Lysate

BCG aliquots were resuspended in 0.75ml of lysis buffer (50mM Tris pH 8.0, 10% glycerol, 0.1% Triton X-100, 1mM PMSF, DNase 3U+2mM MgCl final conc) followed by an incubation period at 30ºC for 15 minutes to facilitate optimal cellular permeabilization. Subsequently, the sonication process was employed, involving three pulses of 30 seconds each for every sample until visual clarity was achieved. Post-sonication, the lysate underwent centrifugation at 12,000 rpm for 20 minutes at 4ºC, ensuring the separation of cellular debris. The protein concentration of the resulting lysate was quantified using the Bradford assay. A concentration of 100 µg/ml was employed as the stock concentration for the aliquots, ensuring a standardized and controlled application for subsequent analyses.

### Flow Cytometry Data Analysis

Isolated spleen cells were transferred to pre-warmed R10 media and washed. Cells were then resuspended at 2-3 million cells per ml in R10 media and stimulated in FACs tubes with *Mycobacterium bovis* antigens or HIV Gag antigens (final concentration of 10μg/mL) in the presence of Golgi-Plug (Cat No: 51-2301KZ, BD Biosciences) and Golgi-Stop(Cat No: 51-2091KZ, BD Biosciences), at a final concentration of 10 μg/mL; for 6h in a 37 °C and 5% CO2 incubator. Samples were stained with Fixability Viability Dye (Zombie NIR, Cat No: 423105 Biolegend) and incubated for 20 mins. Then cells were washed and followed by adding of surface antibody cocktail. After incubating for 30 mins cells were washed and fixed with fixation buffer (Cat No:420801 Biolegend) for 30 mins and permeabilized with 1 × Intracellular Staining Permeabilization Wash Buffer (Cat No: 421002 Biolegend). The optimized concentrations of directly conjugated primary antibodies for intracellular antigen detection were added to samples and incubated for 30 minutes at 4 °C in the dark. Every washing stage was carried out at 700xg for five minutes at 4 °C. All antibodies were previously titrated to determine the optimal concentration (Supplementary Table 1). Finally, after washing and filtering cells through strainer capped FACs tubes, samples were acquired on a Cytek Aurora flow cytometer and analyzed using FlowJo version 10.9.0.

FCS files were exported from SpectralfFo and imported into Flowjo for subsequent analysis. We obtained “fluorescence minus 1” controls (cells stained with all fluorochromes used in the experiment except 1) for each marker prior to spectral unmixing (Supplementary figures 1A-B).

### Statistics

GraphPad Prism 10 was used to compare the means between groups, using the Mann-Whitney non-parametric t-test (two tailed). One asterisk denotes a p-value of less than 0.05, and p-values are indicated in the figure legends and figure images.

## Results

### Multi-Dimensional Flow Cytometry Analysis of Splenic NK Cell Responses to BCG Immunization in C57BL/6 Mice

In this study, we explored the impact of BCG vaccination on NK cell responses in C57BL/6 mice. Four mice were immunized with live BCG Pasteur, while controls received PBS (Figure 1A). After 18 days, spleens were harvested, and single-cell suspensions were analyzed using a 20-color flow cytometry panel (Supplementary Table 1). FlowSOM and Uniform Manifold Approximation and Projection (UMAP) techniques were utilized for visualization and clustering. CD45+CD3-CD19-NK1.1+ cells were analyzed, revealing twenty distinct NK cell clusters (Figure 1B). Heatmap analysis annotated different NK cell subsets based on receptor expression intensity (Figure 1C). Statistical analysis compared vaccinated and control groups, identifying five clusters with increased and three with decreased expression in vaccinated mice (Figure 1C, and Supplementary Figure 2A and Supplementary Figure 2B). Notably, clusters 4 (CD27^dim^PD-1^dim^KLRG1dim), 10 (CCR2+CD27+PD-1^dim^KLRG1++), and 20 (CD27^dim^CD11b+PD-1^dim^KLRG-1+) exhibited higher KLRG1 expression, suggesting phenotypic alterations post-vaccination (Figure 1D). These findings highlight the dynamics of NK cell responses to BCG vaccination, offering insights into potential mechanisms underlying immune modulation.

**Figure 1.**
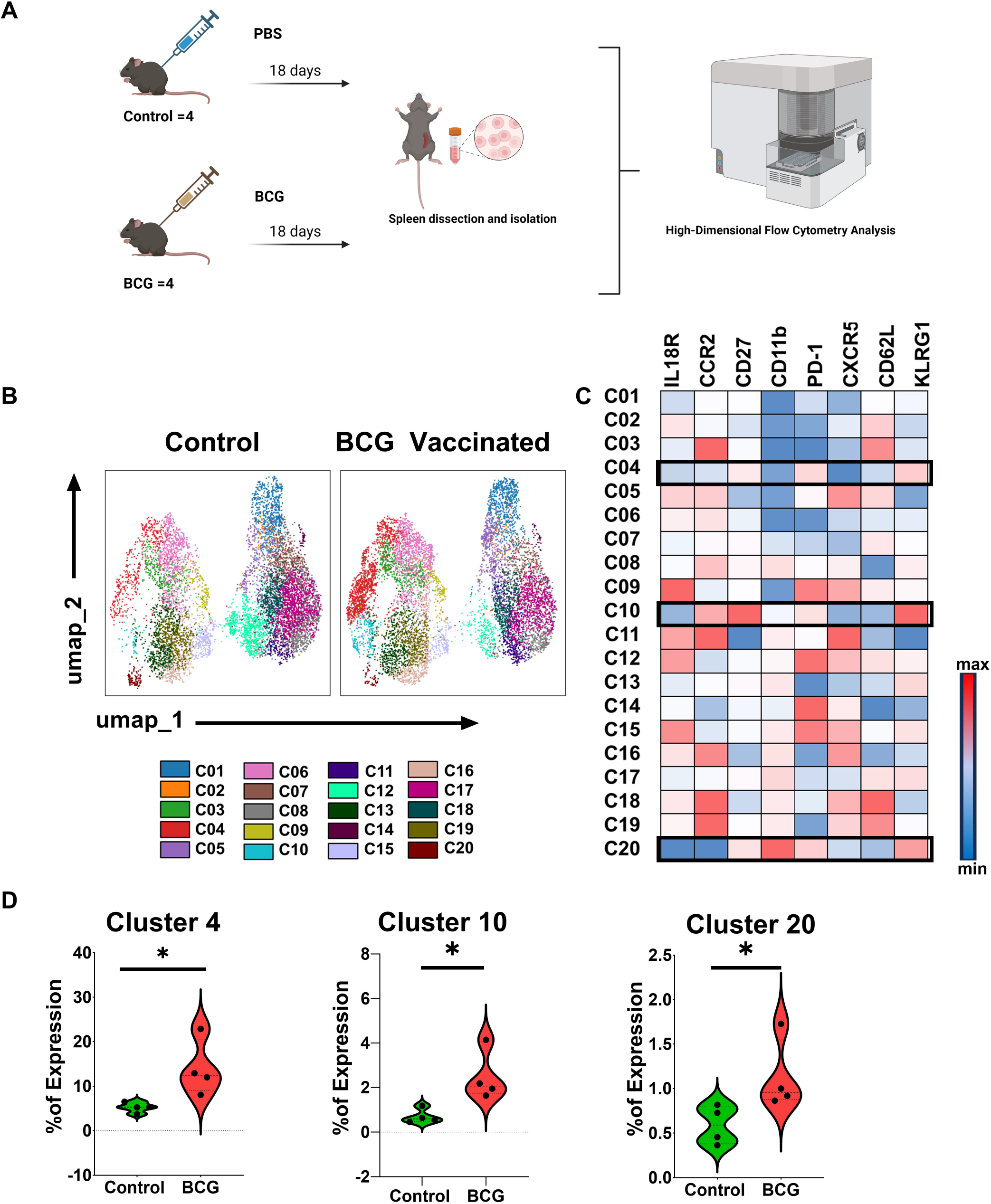
Phenotypic characterization of NK cell populations in spleen in BCG vaccinated and unvaccinated (control) mice. **(A)** Experimental design utilizing C57BL/6 mice. **(B)** FlowSOM-based analysis of viable splenic CD45^+^CD3^-^CD19^-^NK1.1^+^ cells of control and BCG vaccinated groups depicted on a Uniform Manifold Approximation and Projection (UMAP). **(C)** Clusters-by-marker heatmap characterizing the receptor expression patterns of individual clusters. **(D)** Violin plots depicting cluster frequency differences between control and BCG vaccinated samples. The P value was calculated using Wilcoxon rank-sum test; *p<0.05. C, Cluster.

### BCG Immunization Induces the Enrichment of a KLRG1+ NK Cells in Mice

In the high-dimensional analysis, we found higher expression of KLRG1 in NK cells from BCG-immunized mice. KLRG1 serves as a marker of maturation and memory in NK cells(15). Subsequent conventional flow cytometry analysis confirmed a significant expansion of KLRG1+ NK cells in vaccinated groups compared to controls (p<0.05) (Figures 2A, B and Supplementary Figure 2C) as well. However, expression levels of other surface receptors, such as PD-1, CD27, IL18R, CD11b, and CCR2, remained unchanged between the BCG immunized and control groups (Supplementary Figure 2D). Further analysis via t-distributed stochastic embedding (t-SNE) revealed distinctive phenotypic characteristics between KLRG1+ and KLRG1-NK cell subsets within CD45+CD3-CD19-NK1.1+ cells (Figure 2C). However, while CD11b expression was significantly elevated in KLRG1+ NK cells, other markers such as PD-1, CD27, CXCR5 and CCR2 showed no significant differences between the two subsets (Figure 2D-E).

**Figure 2.**
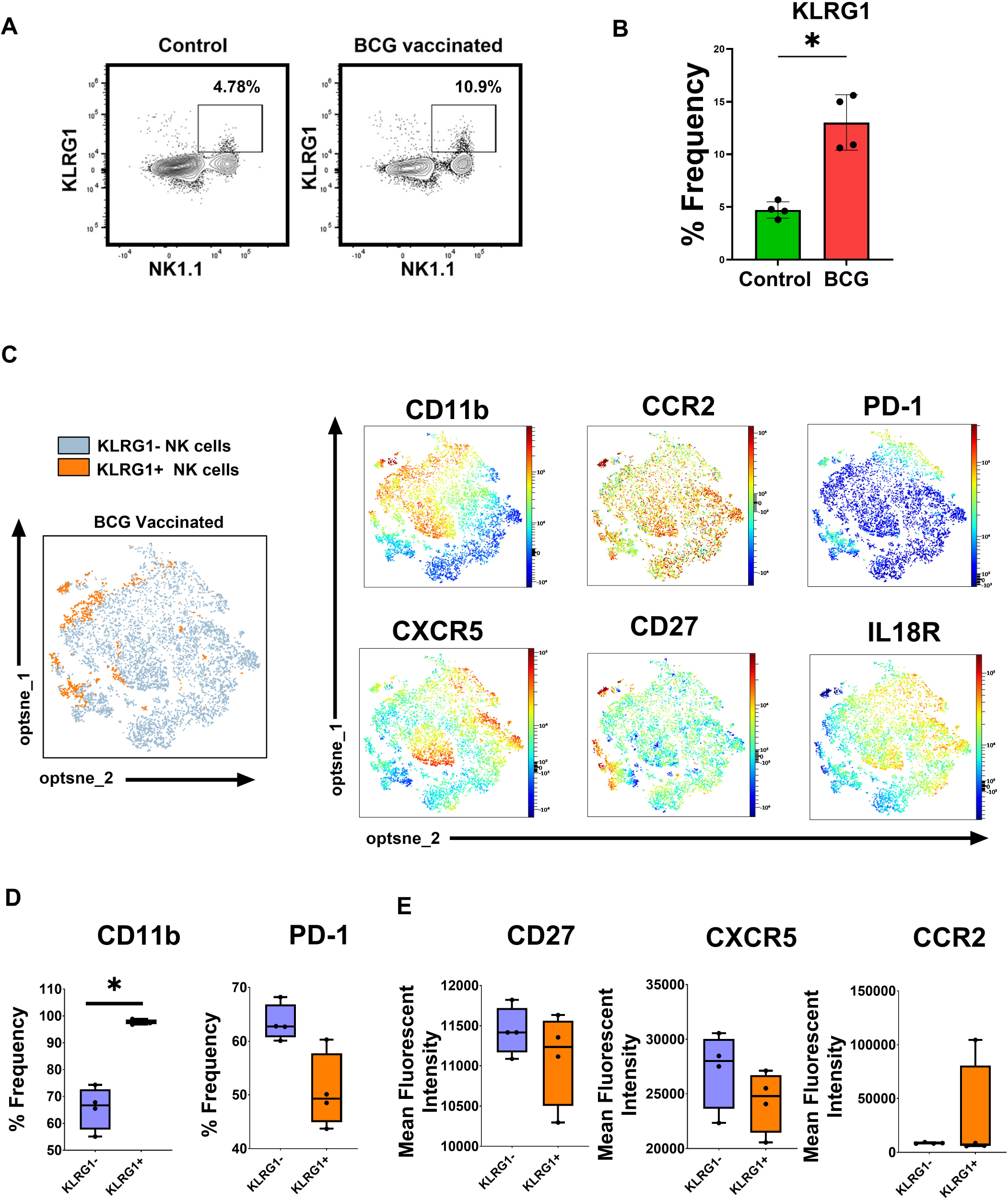
Phenotypic characterization of KLRG1^+^ NK cells in spleen following BCG vaccination. **(A)** Representative contour plots depicting gating strategy for KLRG1^+^ NK cells. **(B)** Box plot illustrating differential proportions of KLRG1^+^ NK cells (among total NK cells) between control and BCG vaccinated mice. **(C)** T-distributed stochastic neighbor embedding (t-SNE) analysis of concatenated samples gated on total viable NK1.1^+^ cells highlighting KLRG1^+^ and KLRG1^-^ NK cell subsets in the BCG vaccinated group. Differential expression of CD11b, CCR2, PD-1, CXCR5, CD27 and IL18R between KLRG1^+^ and KLRG1^-^ NK cell populations visualized by t-SNE. Boxplots comparing **(D)** frequencies and **(E)** Mean Fluorescent Intensities of receptor expression between KLRG1^-^ and KLRG1^+^ NK cells in the BCG vaccinated group. The P value was calculated using Wilcoxon rank-sum test; *p<0.05.

### Memory-like Functional Responses of BCG-Primed NK Cells in Response to Homologous (*Mycobacterium bovis*) Antigens

While there is no universally accepted cytokine correlate of BCG-induced protection against MTB infection, an enhanced capacity of immune cells to produce pro-inflammatory cytokines such as IFNγ, TNFα, etc., may contribute to BCG vaccine efficacy (16). To evaluate BCG-primed NK cell responses to *Mycobacterium bovis* antigens, spleen lymphocytes from vaccinated and control mice were exposed to *Mycobacterium bovis* cell lysate for 6 hours, followed by flow cytometry analysis (Figure 3A). BCG-vaccinated mice exhibited significantly greater increase in production of IFNγ, TNFα, and IL1β in splenic NK cells compared to controls (p<0.05 for all). Granzyme B production in NK cell subsets was also notably higher in vaccinated mice, indicating enhanced cytotoxic potential (p<0.05) (Figure 3B). Polyfunctional NK cells, capable of simultaneous cytokine secretion and granzyme B production, were more prevalent in the BCG-vaccinated group (p<0.05) (Figure 3C and Supplementary Figure 3B). Polyfunctional NK cells, defined by their capacity to simultaneously produce multiple effector molecules, exhibited a heightened response in terms of IFNγ, TNFα, IL1β, and granzyme B production in various combinations (Figure 3C). Following the application of Boolean gating methods to the NK cell subsets(14, 17), percentage of cells producing multiple effector molecules, were significantly elevated in the BCG-vaccinated group (p<0.05). More specifically, we observed significant increases in TNFα^+^ IFNγ^+^ NK cells, TNFα^+^ IL1β^+^ NK cells, IFNγ^+^IL1β^+^ NK cells, TNFα^+^IFNγ^+^IL1β^+^ NK cells and TNFα^+^IL1β^+^granzyme B^+^ NK cells in BCG vaccinated mice after mycobacterial antigen exposure (Supplementary Figure 3B). Next, we compared the cytokine production capacity of KLRG1+ and KLRG1-cell subsets in the BCG-vaccinated group following exposure to *M. bovis* antigens. Interestingly, we found significantly greater increase in IFNγ production in KLRG1+ cells compared to their KLRG1-counterparts. However, we did not observe differences in TNFα, IL-1β, or Granzyme B production between KLRG1+ and KLRG1-subsets in BCG-vaccinated mice upon restimulation with MTB antigens (Figure 3D). These results indicate that KLRG1+ NK cells in BCG-vaccinated mice exhibit unique memory-like responses to mycobacterial antigens.

**Figure 3.**
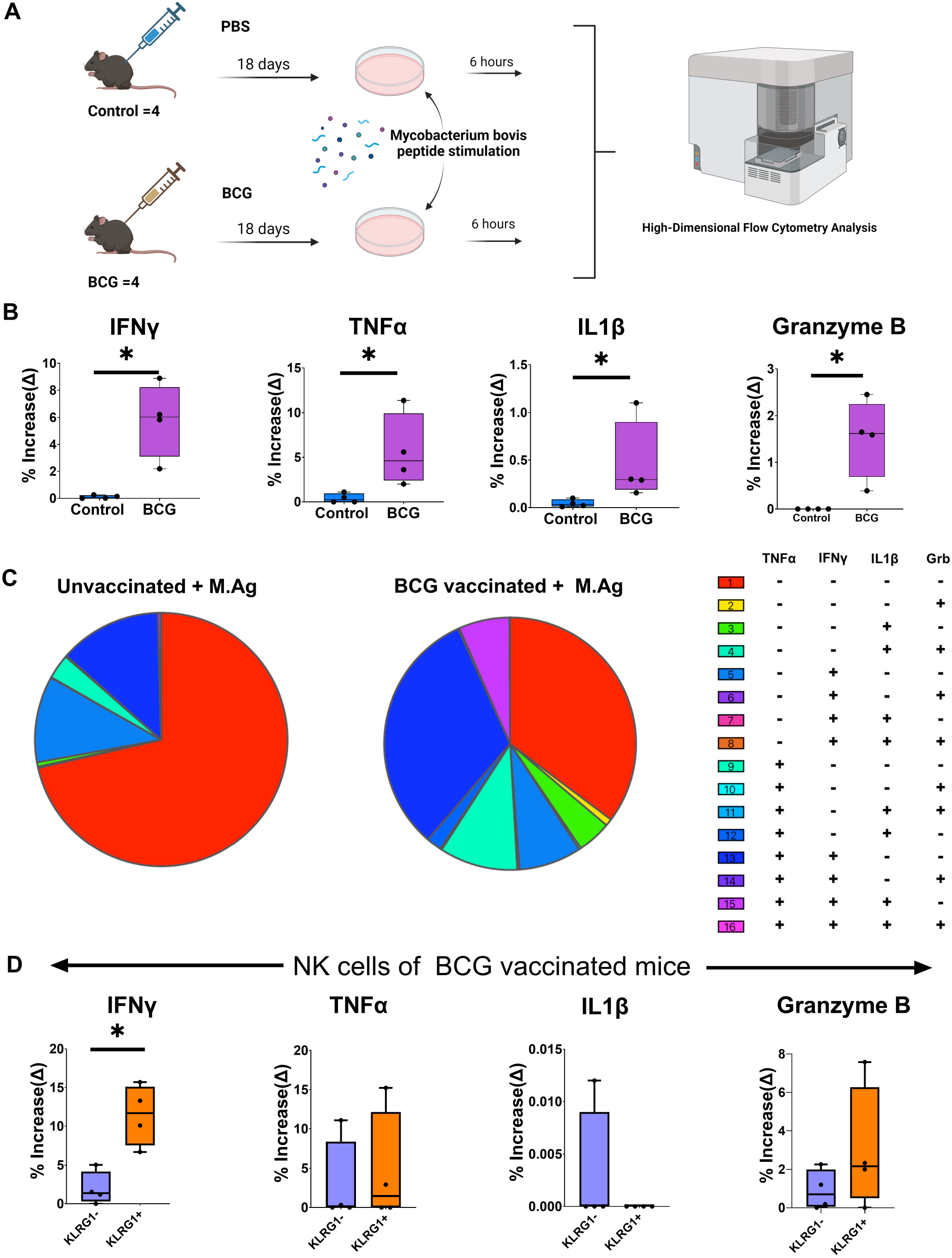
BCG-primed NK cells show enhanced functionality when challenged with mycobacterial antigen. **(A)** Experimental design depicting the in vitro protocol for exposure of splenocytes with *Mycobacterium bovis* antigen. **(B)** Boxplots comparing the increase in expression (Δ values) of IFNγ, TNFα, IL1β and granzyme B (GrB) by NK cells in control and BCG vaccinated mice following *Mycobacterium bovis* antigen exposure. **(C)** Boolean analysis of NK cell polyfunctionality for IFNγ, TNFα, IL1β and GrB production following *Mycobacterium bovis* antigen exposure visualized by SPICE software (v.5.22). **(D)** Boxplots comparing increases (Δ values) in cytokine and GrB production between KLRG1^-^ and KLRG1^+^ NK cells in BCG vaccinated group following *Mycobacterium bovis* antigen exposure. The P value was calculated using Wilcoxon rank-sum test; *p<0.05. Granzyme B,GrB.

### BCG Immunization Induced KLRG1+ NK Cells Demonstrate Enhanced IFNγ Production when Stimulated with Heterologous (HIV gag) Antigen

To study the impact of prior BCG vaccination on NK cell responses to HIV antigens we carried out an in vitro experiment where BCG-primed and control splenic immune cells were exposed to HIV Gag pooled peptides (Figure 4A). BCG-vaccinated and control groups’ splenic cells were stimulated with HIV gag peptides for 6 hours. NK1.1+ cells from BCG group showed no significant changes in levels of IFNγ, TNFα, IL-1β, or Granzyme B production (Figure 4B) or their polyfunctionality compared to controls (Figure 4C).

**Figure 4.**
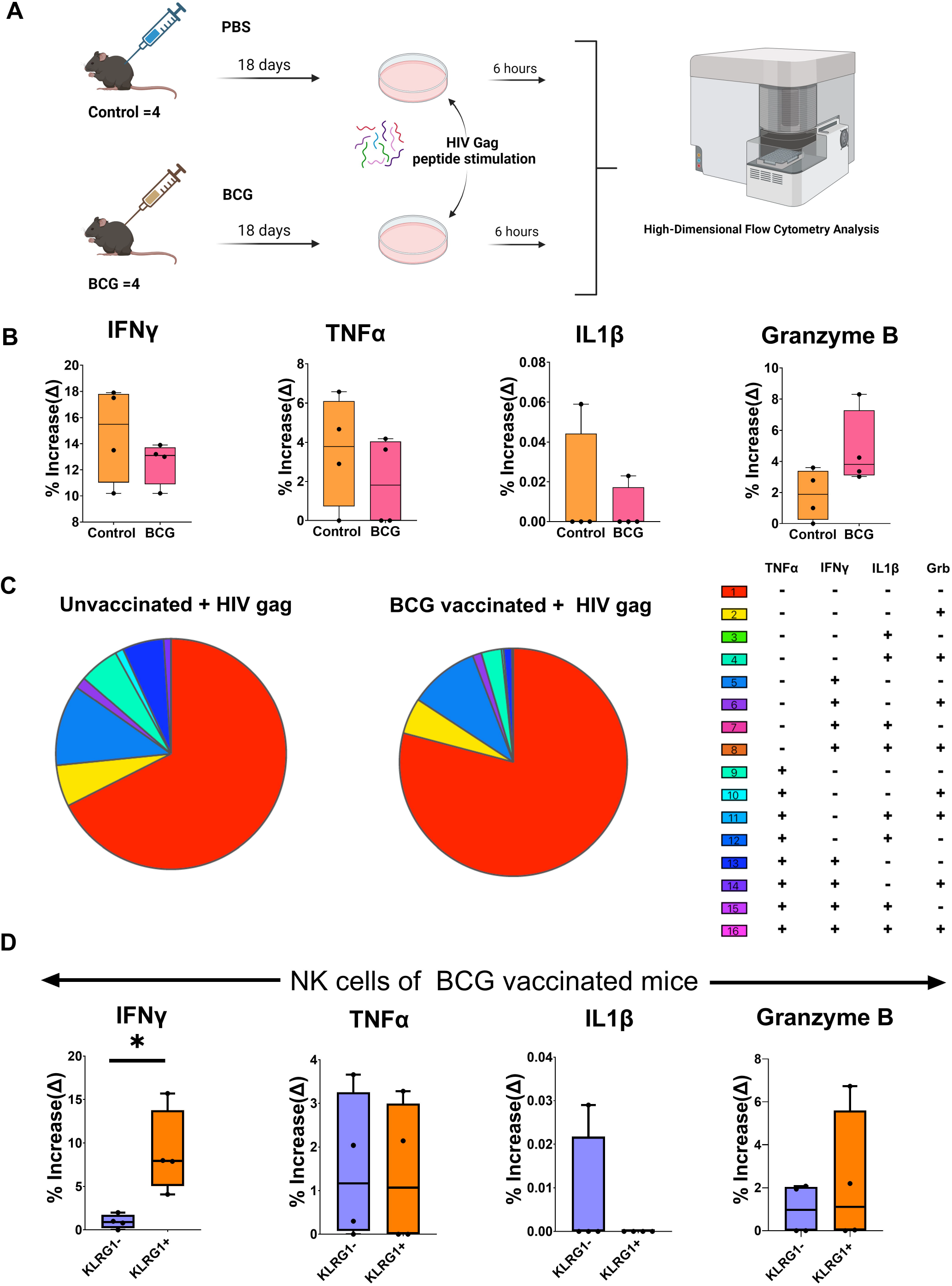
BCG primed KLRG1+ NK cells show enhanced IFNγ production upon exposure to HIV peptides. **(A)** Experimental design depicting the in vitro protocol for exposure of splenocytes with HIV Gag peptides. (B) Boxplots comparing increase in expression (Δ values) of IFNγ, TNFα, IL1β and granzyme B (GrB) by NK cells in control and BCG vaccinated mice following HIV antigen exposure. **(C)** Boolean analysis of NK cell polyfunctionality for IFNγ, TNFα, IL1β and GrB production following *HIV Gag* antigen exposure visualized by SPICE software (v.5.22) **(D)** Boxplots comparing increases (Δ values) in cytokine and GrB production between KLRG1^-^ and KLRG1^+^ NK cells in BCG vaccinated group following HIV antigen exposure. The P value was calculated using Wilcoxon rank-sum test; *p<0.05.

Next, we focused on the functional capability of the expanded KLRG1+ NK cell subset following BCG vaccination, upon exposure to HIV gag peptides. KLRG1+ NK cells from BCG-vaccinated mice demonstrated greater increases in IFNγ production compared to the corresponding KLRG1-NK cells post-stimulation (p<0.05) (Figure 4D). No other significant changes to effector molecules evaluated were affected in KLRG1+ NK cells compared to KLRG1-NK cells. These results show that BCG vaccine-induced KLRG1+ NK cells exhibit heterologous memory-like responses upon exposure to HIV antigens.

## Discussion

In our present study, we show that BCG vaccination leads to the expansion of unique NK cell phenotypes, specifically KLRG1+ NK cells, which exhibit memory-like properties to homologous (*Mycobacterium*) and heterologous (HIV gag) antigens. These cells display an enhanced ability to produce IFNγ upon re-exposure to mycobacterial antigens compared to BCG-exposed KLRG1-NK cells. Additionally, we found that BCG vaccination of mice enhances the production of proinflammatory cytokines and granzyme B by splenic NK cells while increasing their polyfunctionality upon exposure to mycobacterial antigens. However, these NK cells did not show enhanced cytokine production in response to HIV gag peptides.

Several studies have demonstrated that NK cells possess adaptive memory-like properties. The functional reprogramming of these cells reflects in their increased capacity to produce IFNγ and enhanced cytotoxicity upon re-exposure to the training stimulus (12). In this study we provide an additional dimension to this functional enhancement by demonstrating a significant increase in polyfunctionality of BCG-primed NK cells upon stimulation with mycobacterial antigens. Here, we specifically noted expansion of several different subsets of NK cells expressing IFNγ along with one or more cytokines and/or granzyme B suggesting that this may likely be the result of BCG induced functional reprogramming of NK cells. Our study also demonstrated that BCG vaccination expands specific phenotypes of NK cells characterized by increased expression of specific surface receptors, particularly KLRG1. This finding is consistent with previous studies that have reported BCG-induced expansion of subsets of KLRG1+ NK cells (18). Moreover, our observations of increased CD11b expression in these KLRG1+ NK cells, compared to KLRG1-cells, suggest that these cells may have a greater capacity for cytokine secretion and cytolytic activity (19). Increased expression of KLRG1 has been noted on antigen-experienced cells, and in our study this priming is likely occurred through BCG vaccination(20, 21). Our data further support previous studies suggesting that KLRG1 defines subsets of memory-like NK cells with protective features against *Mycobacterium tuberculosis*. We found that the expanded KLRG1+ NK cells, primed by BCG possess an increased capacity to produce IFNγ upon exposure to mycobacterial antigens, compared to their KLRG1^-^ counterparts(18).

Furthermore, we demonstrated that BCG vaccination induces non-specific priming of KLRG1^+^ NK cells, as evidenced by their increased production of IFNγ in response to unrelated stimuli such as HIV gag. The significance of this expanded KLRG1+ NK phenotype as a subset with non-specific memory-like properties was further highlighted when exposure to HIV gag peptides failed to induce an overall increase in IFNγ (and/or other tested cytokines and granzyme B) or polyfunctionality in total NK cells of BCG vaccinated mice.

Previous studies have indeed identified unique subsets of NK cells with memory-like properties that undergo expansion in response to priming by various stimuli that generate trained immunity. For instance, In human cytomegalovirus (CMV) seropositive individuals, NKG2C^hi^ NK cells were found to be expanded and were considered memory-like NK cells; they specifically showed epigenetic remodeling of the conserved non-coding sequence region in the *IFNG* locus, facilitating *IFNG* transcription upon signaling from activating receptors(13). Similarly, studies on murine CMV infection have identified Ly49H as a marker of NK cell memory(12).

However, whether epigenetic changes underline the BCG induced increased expression of KLRG1 in our subset of memory-like NK cells, demonstrating enhanced IFNγ production to related and unrelated viral (i.e., HIV gag) antigens remains to be investigated. Deep analysis of the epigenetic profiles of these NK cell subsets is warranted to address this question.

While our study has several limitations, including a small sample size, an analysis incorporating only a limited number of NK cell surface markers and cytokines, and an analysis restricted to splenic tissue, we present compelling evidence that BCG vaccination expands phenotypic subsets of NK cells with memory like features. This is indicated by the increased IFNγ production by these cells upon challenging with related bacterial antigens and unrelated viral antigens such as HIV gag peptides. Despite limitations, our findings may have significant implications, particularly in expanding the benefits of existing vaccines. Our data justifies further studies aimed at exploiting the advantages of trained immunity through innovative vaccine design. This calls for the careful selection of vaccine antigens and adjuvants that can initiate innate immune memory, potentially leading to protection against multiple pathogens.

## Author Contributions

M.G performed experiments, data analysis and, wrote the manuscript. M.A wrote and edited the manuscript. W.M aided in literature survey and figure editing. T.D, edited the manuscript. N.L designed the study, supervised data analysis and wrote the manuscript.

## Acknowledgments

The authors gratefully acknowledge and thank Richard Robinson for providing the *Mycobacterium bovis* BCG Pasteur stocks and help with the experiments with mice.

