## Supplemental Figures for "BCG immunization induced KLRG1+ NK cells show memory-like responses to mycobacterial and HIV antigens"

A

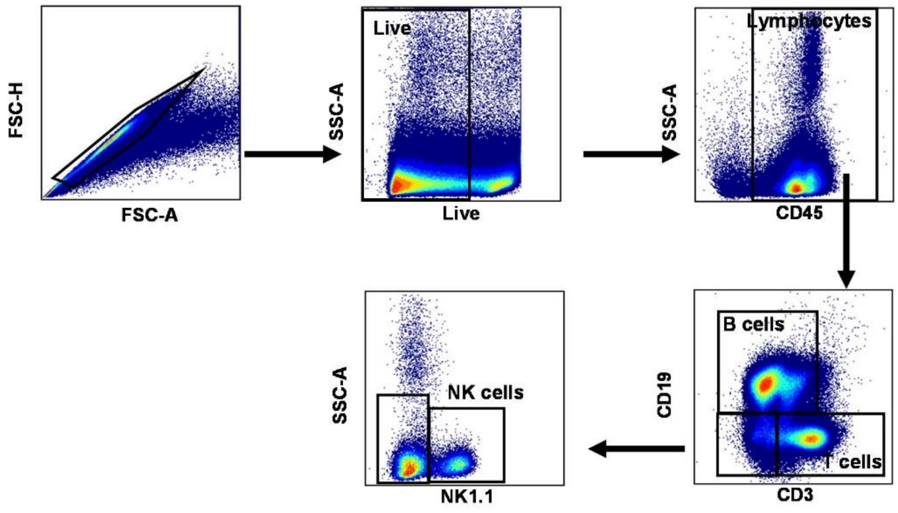


B

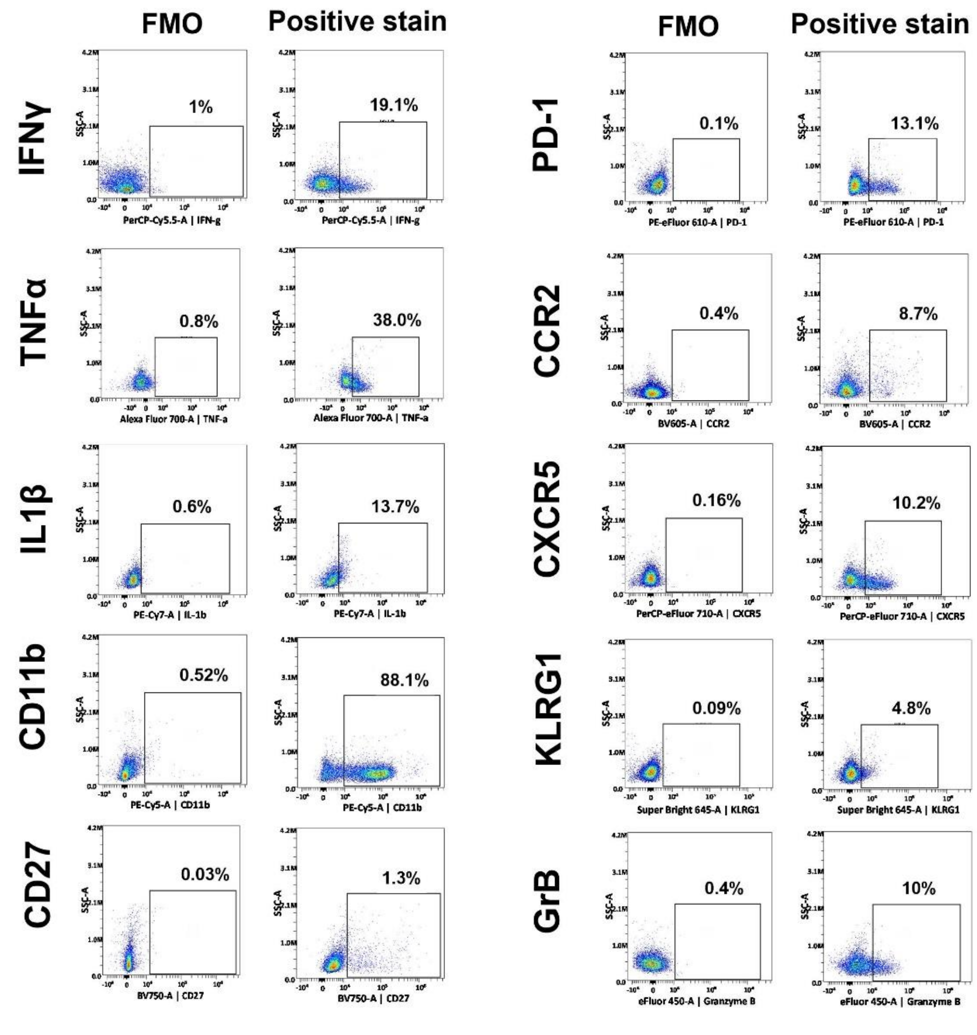


Supplementary 1 Figure. (A) Representative pseudo color plots depicting flow cytometric gating strategy used to define immune cell populations in mouse splenocytes. (B) Representative pseudo color plots illustrating fluorescent minus one controls for receptor expression. GrB, Granzyme B.

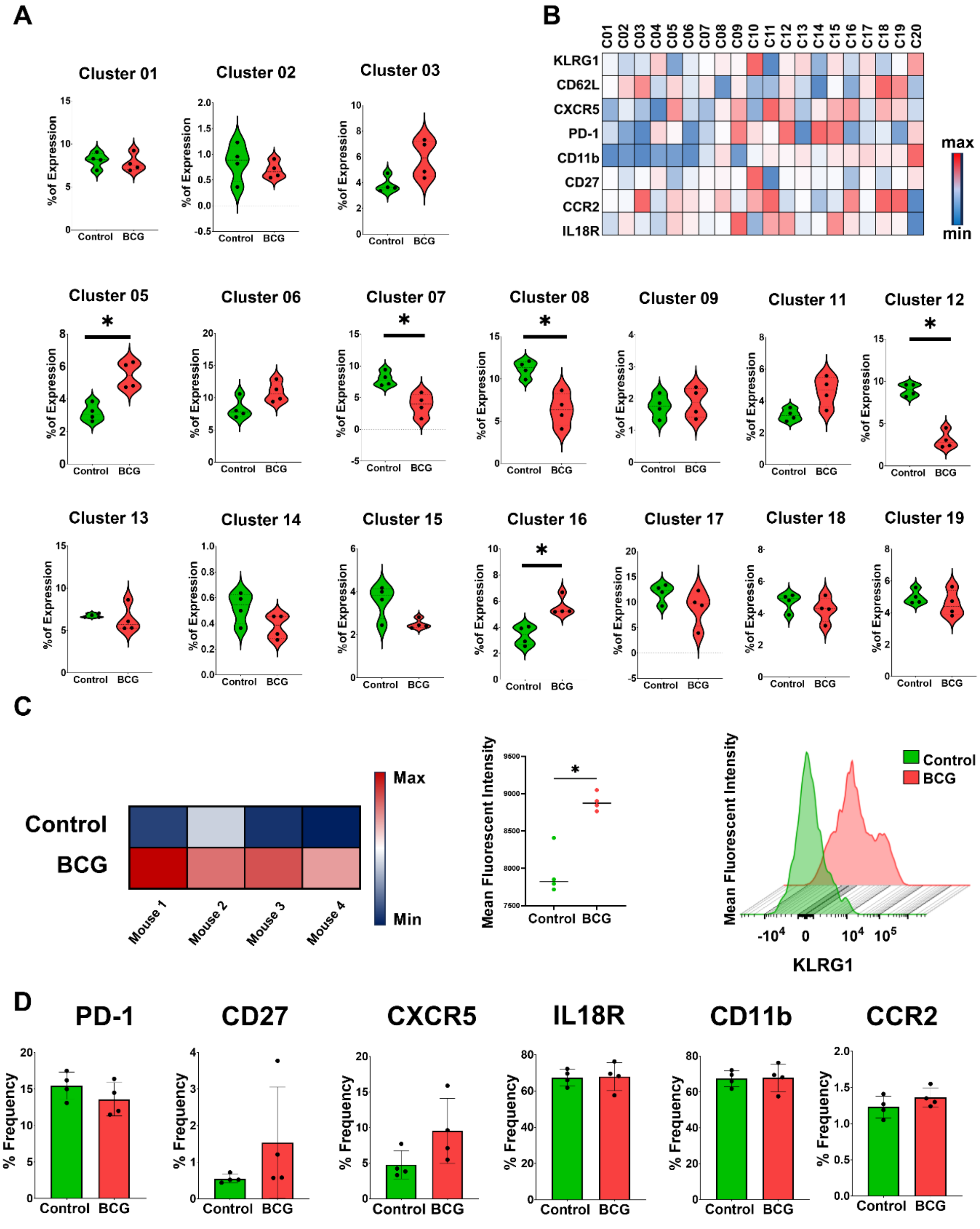


Supplementary 2 Figure. (A) Violin plots depicting NK cell cluster frequencies of Control vs BCG vaccinated samples. (B) Clusters-by-marker heatmap characterizing the receptor expression patterns of individual clusters. (C) Heatmap, dot plot and (representative) histogram comparing Mean Fluorescence Intensity of KLRG1 expression for all eight animals in the study. (D) Bar plots depicting receptor expression levels for PD-1, CD27, CXCR5, CD11b, IL18R, and CCR2 in NK cells of Control and BCG vaccinated mice.

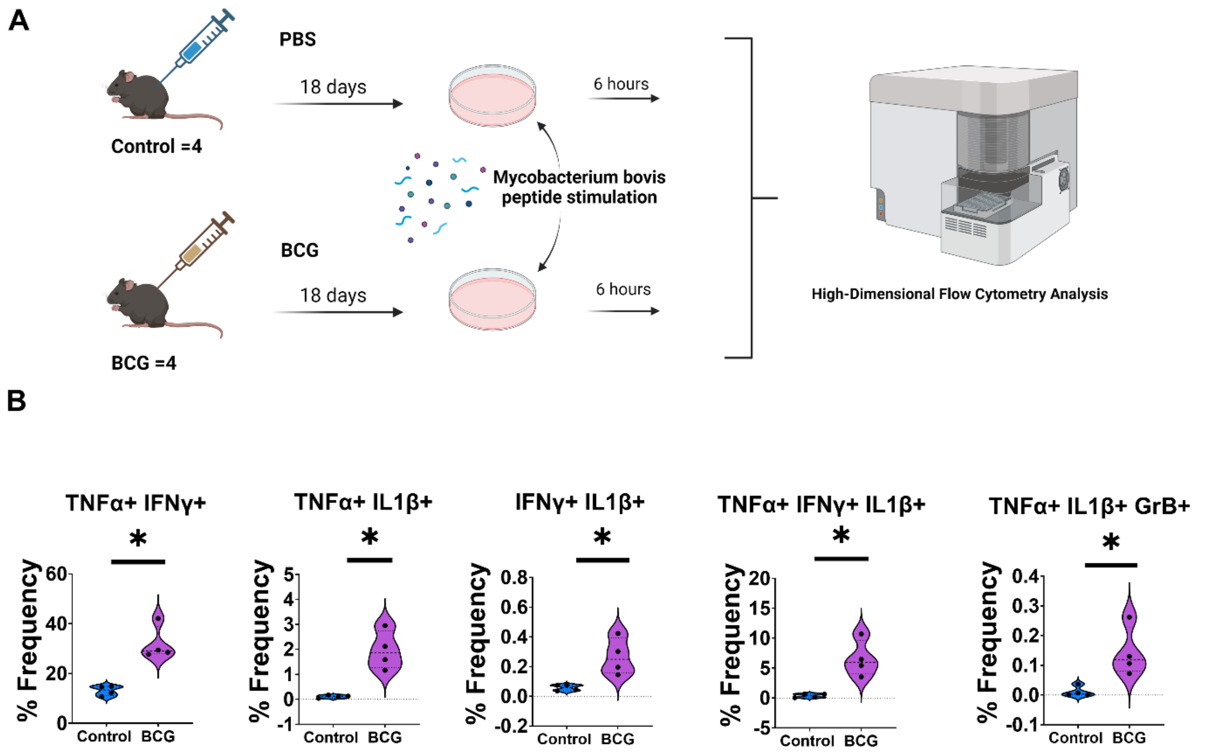


Supplementary 3 Figure. (A) Experimental design depicting the in vitro protocol for exposure of splenocytes with *Mycobacterium bovis* antigen. (B) Violin plots illustrate the significant differences in proportions of polyfunctional NK cell subsets between control and BCG vaccinated groups, based on production of IFNγ, TNFα, IL1β, and Granzyme B following exposure of splenocytes with *Mycobacterium bovis* antigen. *$p<0.05$. Granzyme B,GrB

Supplementary Table 1: List of antibodies for Flow Cytometric analyses

| Fluorochrome | Specificity | Clone | Company | Cat.No | Concentration |
|---|---|---|---|---|---|
| FITC | CD3 | 17A2 | Biolegend | 100203 | 1:100 |
| PE- eFluor 610 | PD-1 | J43 | Invitrogen | 61-9985-82 | 1:50 |
| PE-Cy5 | CD11b | M1/70 | Invitrogen | 15-0112-83 | 1:50 |
| PerCP-Cy5.5 | IFNγ | XMG1.2 | Biolegend | 505821 | 1:50 |
| PerCP-eFluor 710 | CXCR5 | SPRCL5 | Invitrogen | 46-7185-82 | 1:50 |
| PE-Cy7 | IL1β | NJTEN3 | eBiosciences | 25-7114-82 | 1:25 |
| Alexa Fluor 647 | IL18R | A17071D | Biolegend | 157907 | 1:50 |
| Alexa Fluor 700 | TNFα | MP6-XT22 | Biolegend | 506338 | 1:50 |
| Brilliant Violet 421 | CD45 | 30-F11 | Biolegend | 103134 | 1:100 |
| eFluor 450 | Granzyme B | NGZB | eBiosciences | 48-8898-82 | 1:50 |
| Brillant Violet 605 | CCR2 | SA203G11 | Biolegend | 150615 | 1:50 |
| Super Bright 645 | KLRG1 | 2F1 | Invitrogen | 64-5893-82 | 1:50 |
| Super Bright 436 | CD62L | MEL-14 | Invitrogen | 62-0621-82 | 1:50 |
| Brilliant Violet 711 | CD19 | 6D5 | Biolegend | 115555 | 1:50 |
| Brilliant Violet 750 | CD27 | LG.3A10 | BD | 747399 | 1:50 |
| Brilliant Violet 785 | NK1.1 | PK136 | Biolegend | 108749 | 1:50 |
